# A Bayesian approach to inferring chemical signal timing and amplitude in a temporal logic gate using the cell population distributional response

**DOI:** 10.1101/087379

**Authors:** Ania A. Baetica, Thomas A. Catanach, Victoria Hsiao, Richard M. Murray, James L. Beck

## 1 Introduction

Stochastic gene expression poses an important challenge for engineering robust behaviors in a heterogeneous cell population [1]. Cells address this challenge using the distribution of cellular responses during some gene regulation and differentiation processes [2]. Similarly, the temporal logic gate design in Hsiao et al. [3] considers the distribution of responses across a cell population. The design employs integrases Bxb1 and TP901-1 [4] to engineer an *E. coli* strain with four DNA states that record the temporal order of two chemical signals. Hsiao et al. [3] also use the heterogeneous cell population response to infer the timing and duration of the two chemical signals for a small set of events. Our work uses the temporal logic gate circuit to address the problem of extracting information about events that were recorded in the distributional response of a cell population. We use the heterogeneous cell population response to infer whether any event has occurred or not and also to infer its properties such as timing and amplitude.

Bayesian inference provides a natural framework to answer questions about chemical signal occurrence, timing, and amplitude [5]. We develop a probabilistic model based on the temporal logic gate model in [3] that integrates both predictive modeling uncertainty and sampling error statistics. In this way, we incorporate uncertainty in how well our temporal logic gate model captures the cell population and in how well a sample of measured cells represents the entire population. Using our probabilistic model and cell population measurements taken every five minutes on simulated data, we ask how likely it was to observe the data for parameter values that describe square-shaped inducer pulses. The likelihood function associated with the probabilistic model answers the question of how likely the data is by comparing the likelihood values for the model where chemical signal pulses are turned off against the model where the pulses are on. Hence, we determine whether an event of chemical induction of integrase expression has occurred or not.

Using Markov Chain Monte Carlo [6], we sample the posterior distribution of pulse parameters and then estimate the posterior probability of the two chemical signal events. We implement this method and obtain accurate results for detecting chemical inducer pulse timing, length, and amplitude. We can detect and identify chemical inducer pulses as short as half an hour, as well as all pulse amplitudes that fall under biologically relevant conditions.

Using the Bayesian framework to solve our problem enables us to obtain distributions over chemical signal occurrence, timing, and amplitude, as well as to test the limits of chemical input identifiability. There are alternative methods for the problems posed in this work. The first problem of detecting an event falls under the broad field of anomaly and change point detection. Many approaches within this field deal with the problem of using models or approximations to quantify the typical behavior of a system and then setting a threshold to determine whether the signal is within this typical set of behaviors. There are data driven methods like clustering methods or spectral methods that do not need a physical model of the system, but use data to determine a typical set of behaviors [7]. Alternatively, when a physical model is known and computationally tractable, a hypothesis testing framework can be employed. If a set of possible models of the event is also known, a likelihood ratio test is often used to determine whether to accept or reject the hypothesis [8]. However, the methodology of Bayesian model selection enables us to better handle uncertainty within the models, to avoid setting detection thresholds that are not based on probabilities, and to avoid overfitting [5, 9, 10].

The second problem of identifying the parameters of the chemical inducer pulse is often approximated by solving a maximum likelihood or maximum posterior estimation problem to find the best set of parameters that describe the data [11, 12]. While the optimization problem can be nonconvex, several methods have been developed to find solutions for these problems such as Expectation-Maximization [12]. Maximum likelihood methods can also be integrated into the change point detection framework [8]. However, using MCMC to sample the posterior distribution of likely parameters enables us to make a more robust estimate about the set of possible pulse parameters. This also enables us to detect when the pulse is weak since this corresponds to the posterior being very broad and close to unidentifiable.

Our paper is organized as follows. In Section 2, we introduce the temporal logic gate circuit and we simulate its behavior over a heterogeneous cell population. In Section 3, we set up the Bayesian inference framework and the Markov Chain Monte Carlo methods. We illustrate the results of applying the Bayesian framework to the problem of inferring chemical inducer properties in Section 4. We discuss our results and future work in the conclusion section.

## 2 The temporal logic gate circuit

### 2.1 The temporal logic gate function

Cellular processes are subject to stochastic fluctuations, particularly at low molecule numbers [1]. This poses an important challenge for engineering robust behaviors in a heterogeneous cell population since we often are not able to design for a homogeneous response. Cells address this challenge by operating on distributions of cellular responses during noisy processes such as probabilistic differentiation [2]. Similarly, the temporal logic gate design in [3] operates on the distribution of cellular responses across a population. The two-input temporal logic gate uses integrases Bxb1 and TP901-1 to engineer an *E. coli* strain with four possible DNA states that record the temporal order of chemical inputs. Using the heterogeneous response of the *E. coli* population, Hsiao et al. [3] infer and record the order of chemical inputs. They also use the heterogeneous population response to infer the timing and duration of the two chemical inputs for a discrete set of events.

The design of the integrase temporal logic circuit uses serine integrases TP901-1 (int A) and Bxb1 (int B) to flip the DNA between their recognition sites, as in Figure 1. Each cell in the population can be in one of 4 identifiable DNA states: no input (state *S*_0_), only input **a** detected (state *S_a_*), only input **b** detected (state *S_b_*) or input **a** then **b** detected (state *S_ab_*) as illustrated in Table 1. When inducer **b** is detected before inducer **a**, the DNA between the attachment sites is excised, so the cells are in state *S_b_*. Fluorescent proteins mKate2-RFP (RFP) and superfolder-GFP (GFP) are used to read the DNA state of each cell. RFP is produced when the cell is in state *S_a_* and GFP is produced in state *S_ab_*. Once a DNA recombination step has occurred, due to the detection of either input **a** or **b**, it is irreversible and thus recorded in DNA memory. The temporal logic gate circuit is integrated chromosomally into the genome of *E. coli* cells [13]. Hence, we can assume that each cell contains only one copy of the circuit and thus its DNA is in one of the four identifiable states *S*_0_, *S_a_*, *S_b_* or *S_ab_*. For more details on the temporal logic gate circuit, see reference [3].

**Figure 1:**
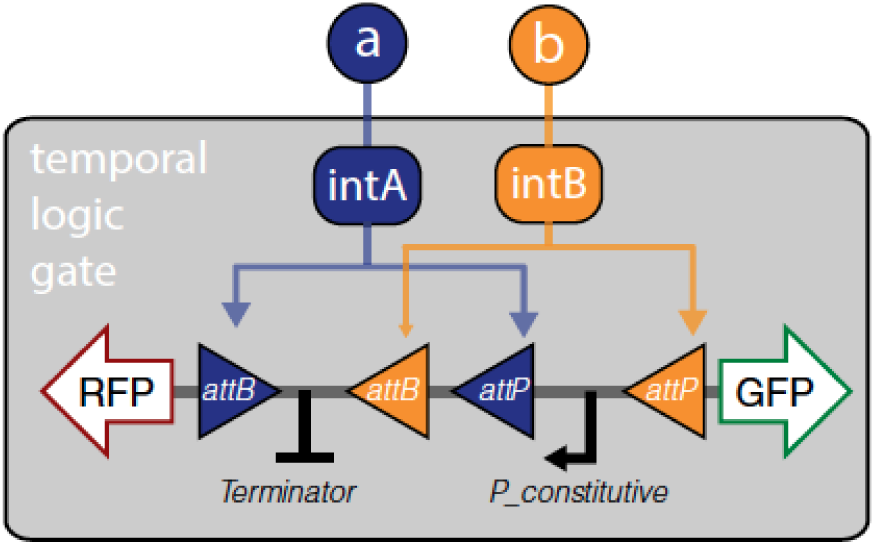
Implementation of the temporal logic gate using a set of two integrases with overlapping attachment sites. Chemical inputs **a** and **b** activate production of integrases intA and intB, which act on a chromosomal DNA cassette. Figure reproduced from reference [3].

**Table 1:**
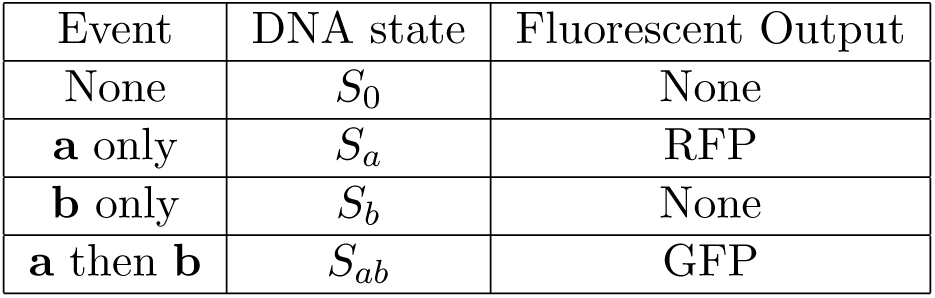
The table describes the inputs, DNA states, and outputs to the temporal logic gate. Table adapted from [3].

### 2.2 Stochastic modeling of the temporal logic gate using the chemical master equation

To capture the stochastic behavior of the temporal logic gate, we model DNA and integrase interactions in each cell using the chemical master equation [14]. Since the target DNA is chromosomally integrated in the *E. coli* genome, we can assume that each *E. coli* cell in the population can be uniquely characterized by the triplet of DNA state and copy numbers of integrases A and B [13].

To build the state space of our Markov chain [15], we follow the supplementary information in [3] and denote the state of a cell as (*S_i_, n_A_, n_B_*), where DNA state *S_i_ ∈* **S** = *{S*_0_, *S_a_, S_b_, S_ab_}* and integrase copy numbers *n_A_, n_B_ ∈* ℕ_*≥*0_. The details of the Markov transitions between states are found in Table 2. The Markov transition rates between states emerge from Figure 2. The chemical master equation (CME) model is described in detail in the supplementary information of [3].

**Table 2:**
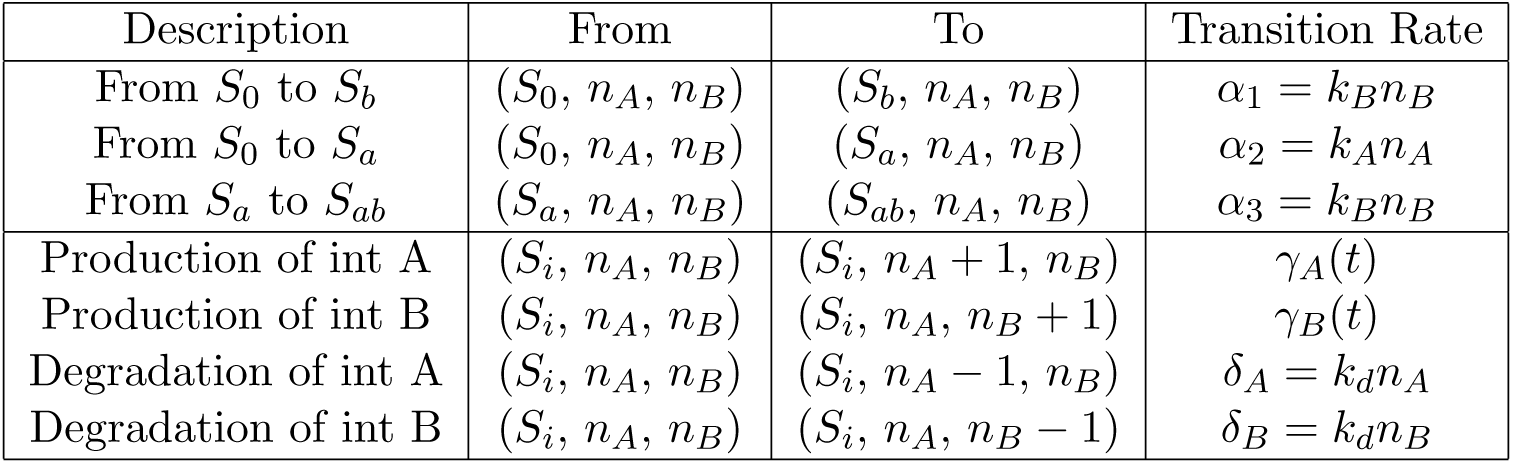
The Markov transition rates between states. *k_A_* and *k_B_* are the DNA flipping rates of integrases A and B. *k_d_* is the degradation rate, while *γ_A_* and *γ_B_* are the production rates of integrases A and B. Table adapted from the supplementary information in [3].

**Figure 2:**
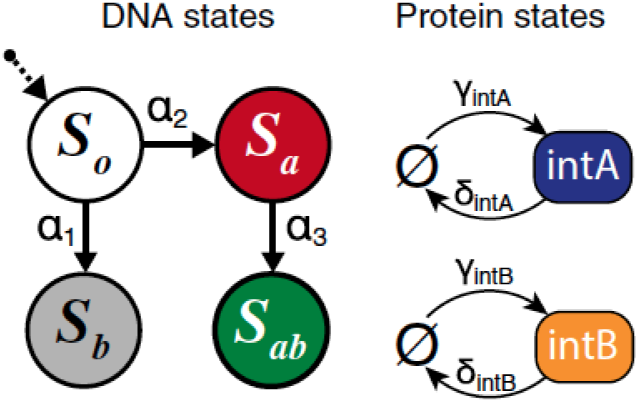
Transitions between DNA states and between protein states. The transition rates are listed above the arrows. Figure reproduced from [3].

The chemical inputs enter the Markov transition rates through the terms *γ_A_*(*t*) and *γ_B_*(*t*) that represent the production of the two integrases. The chemical inputs that we consider are square waves as illustrated in Figure 3. We assume that chemical inputs **a** and **b** are turned off, they turn on, and then they turn back off. This defines our production rates *γ_A_*(*t*) and *γ_B_*(*t*) as

**Figure 3:**
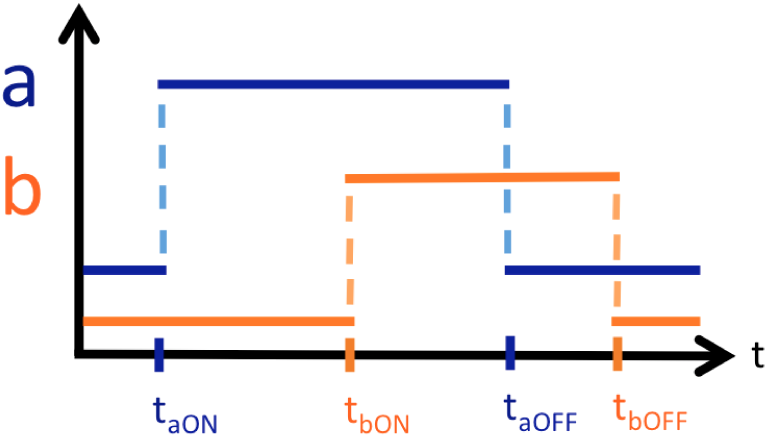
We restrict the chemical inducer inputs **a** and **b** to be square waves. They turn on at times *t*_aON_ and *t*_bON_ and turn off at times *t*_aOFF_ and *t*_bOFF_.

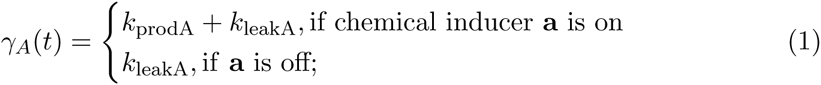

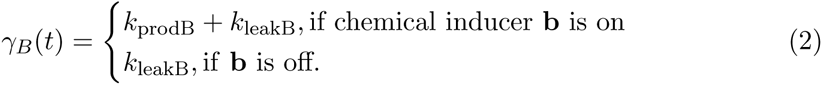

Here *k*_prodA_ and *k*_prodB_ are the production rates of the two integrases, while *k*_leakA_ and *k*_leakB_ represent leakiness.

It is possible to consider chemical input functions that are not square waves, but rather arbitrary continuous functions. In this case, the CME will have a numerical solution, but possibly not an analytic one. Hence, we restrict our chemical inputs to square wave functions.

### 2.3 Solving the chemical master equation model

We simulate the CME model of the temporal logic gate circuit using the finite state projection algorithm (FSP) in [16] as follows. We first transform our three dimensional state space into a one dimensional state vector by iterating over the four DNA states and the pairs of integrase copy numbers. The resulting infinite state space vector is given by:

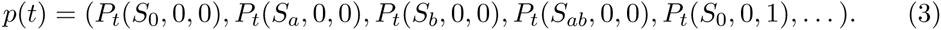

The transition matrix *A* is square and extends to infinity since the state vector has an infinite number of entries. We separate our transition matrix into a sum of a constant matrix and two matrices multiplied by production rates:

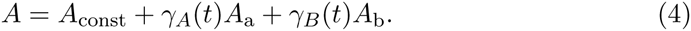

This separation speeds up simulation as there is no need to recompute the matrices *A_a_* and *A_b_* as the chemical input functions *γ_A_* and *γ_B_* vary in time. Moreover, if the promoters are non-leaky and one or both chemical inputs are turned off, then equation (4) further simplifies, thus speeding up computation.

Matrices *A*const, *A*_a_, and *A*_b_ are still infinite in both of their dimensions. We truncate them according to the FSP algorithm to a maximum of 20 copies of integrases A and B in each dimension. The truncation is informed by experimental data in [3]. The total amount of probability lost in the exponential of the transition matrix by this truncation is less than 0.01.

Following the truncation of the transition matrix using FSP, the CME formulation of the temporal logic gate model is

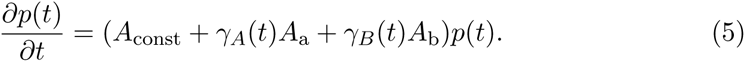

We solve for the probability distribution over the heterogeneous cell population by computing the standard matrix exponential solution

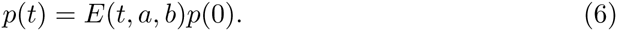

The matrix *E*(*t, a, b*) is the product of exponential matrices according to the ordering and time at which inducers **a** and **b** turn on and off. For example, if inducer **a** turns on instantaneously and inducer **b** turns on at time *t*_bON_ and they both subsequently remain on, then the expression for *E*(*t, a, b*) at time *t* follows from:

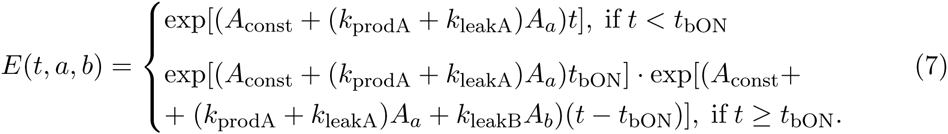

Similar expressions for the matrix *E*(*t, a, b*) can be derived for any combination of chemical inducers **a** and **b**. The result can be then used in equation (6) to derive how the heterogeneous cell population evolves as a function of time by patching together the different solutions at each time interval.

### 2.4 Simulation results for the temporal logic gate circuit model

We simulate the chemical equation model of the temporal logic gate circuit in MATLAB [17]. In Table 3, we record the times when the chemical inducers turn on and off. In Figures 4 and 5, inducer **b** has been on for only 3 hours, so cells in states *S*_0_ and *S_a_* make up most of the population. They have either mostly responded to inducer **a** or they have not seen any of the two chemical inducers. Cells in state *S*_0_ will shift to state *S_b_* and cells in state *S_a_* will shift to state *S_ab_* as they continue to be exposed to chemical inducer **b**.

**Table 3:**
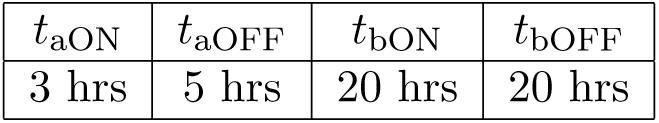
The chemical inducer **a** and **b** turn on and off times for the simulation results in Figures 4 - 7.

**Figure 4:**
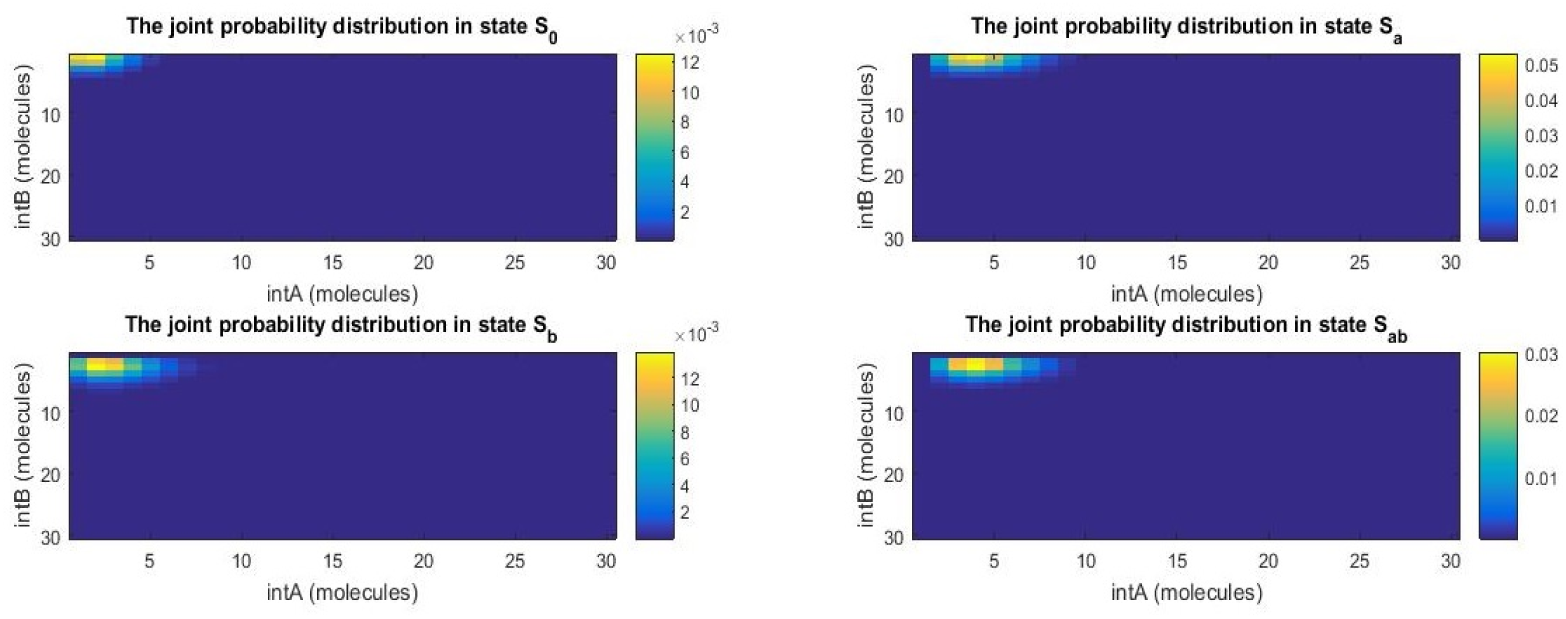
The four panels represent the two-dimensional probability distributions of cells in states *S*_0_, *S_a_*, *S_b_*, and *S_ab_* as functions of integrase copy numbers after 8 hours. Inducer **a** has been turned on at 3 hours and inducer **b** at 5 hours. Most cells have only seen inducer **a** and are in state *S_a_*, although some cells are starting to detect inducer **b**.

**Figure 5:**
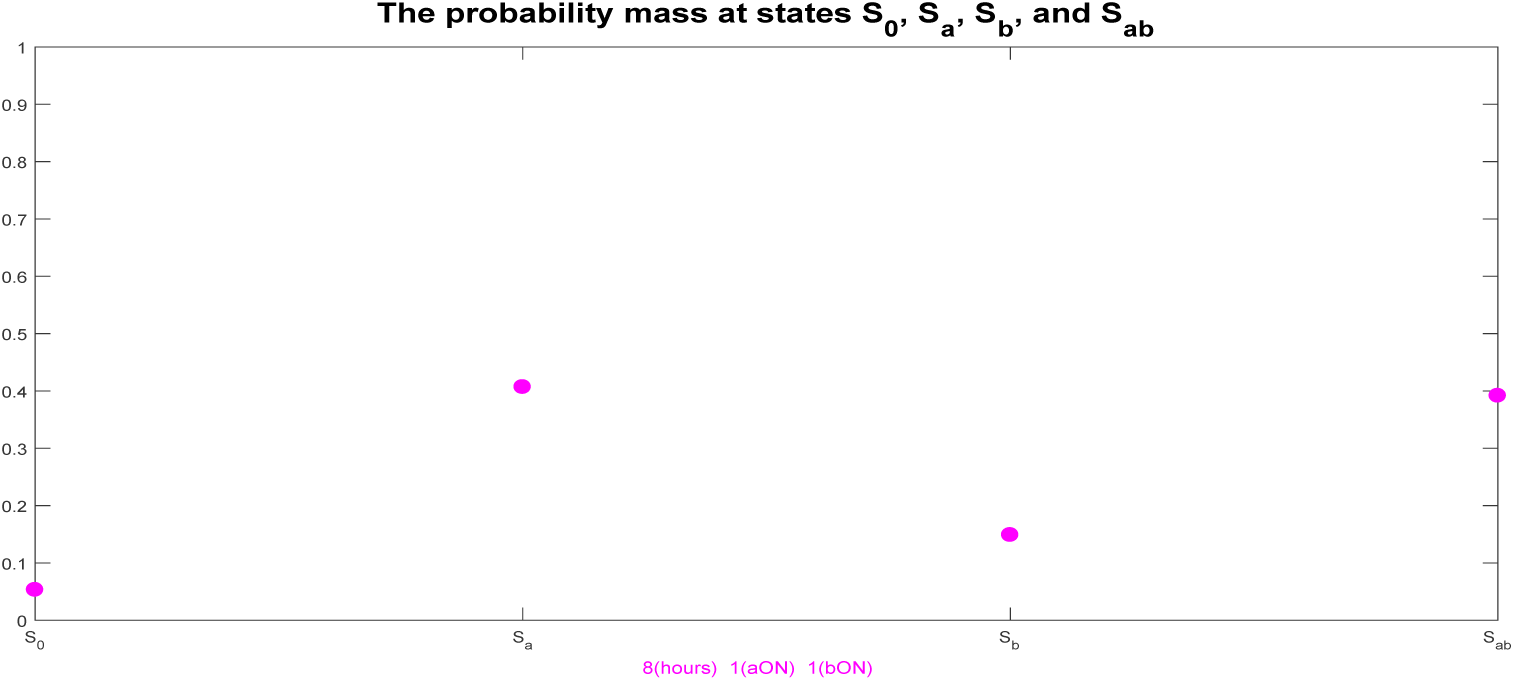
The plot illustrates the fraction of cells in each of the four states. The timer at the bottom indicates that the circuit has been running for 8 hours. The Boolean variables next to input **a** and input **b** show that **a** has been turned on at 3 hours and **b** at 5 hours. Not all cells have responded to chemical inducer **b** yet, as indicated by the fraction of cells in state *S_a_*.

In Figures 6 and 7, both inducers have been on for several hours, so cells have shifted to states *S_b_*, if they have only seen inducer **b**, and *S_ab_*, if they have recorded the “**a** then **b**” event. This population behavior matches the experimental evidence in [3]. The rate reaction parameters have the same values as in the supplemental information of [3]. We use the simulation data in this section for the Bayesian approach to inferring parameter properties that we introduce in Section 3.

**Figure 6:**
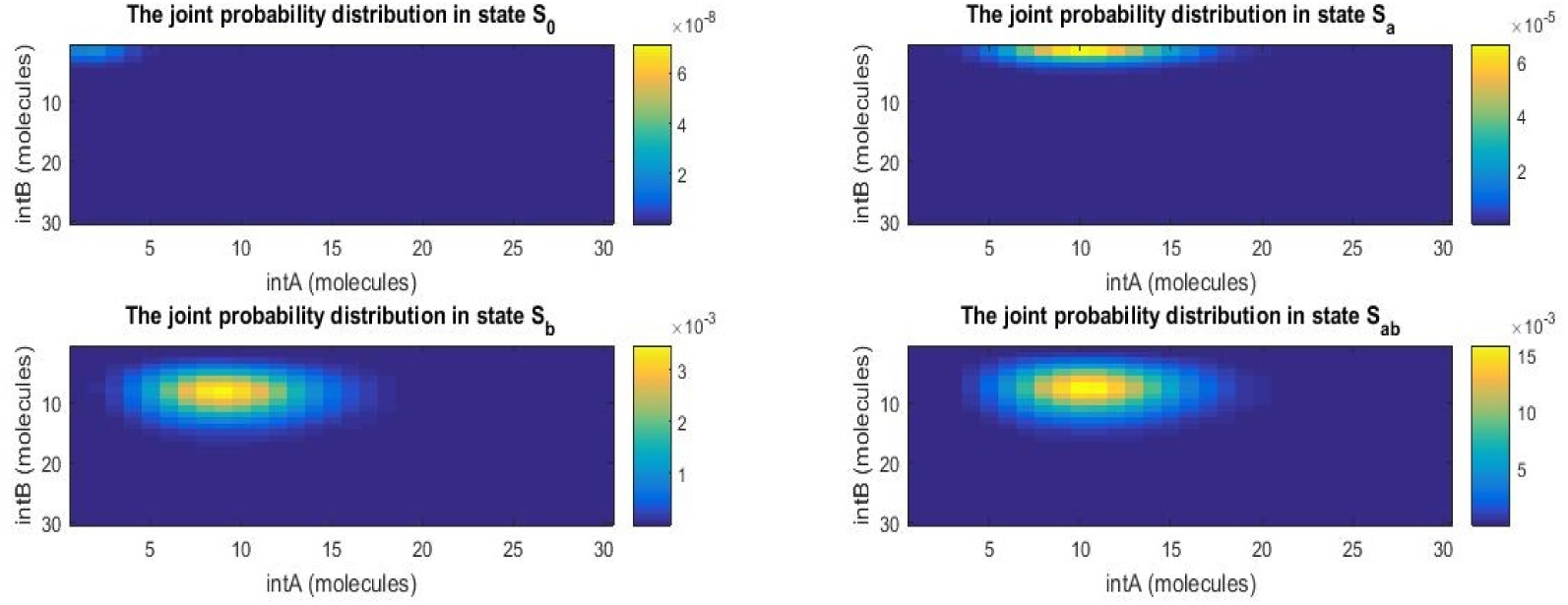
The temporal logic gate circuit has been running for 20 hours and it has reached stationary state. Input **a** has been turned on at 3 hours and input **b** has been turned on at 5 hours; subsequently, they were both on. Most cells are either in state *S_b_* or state *S_ab_*.

**Figure 7:**
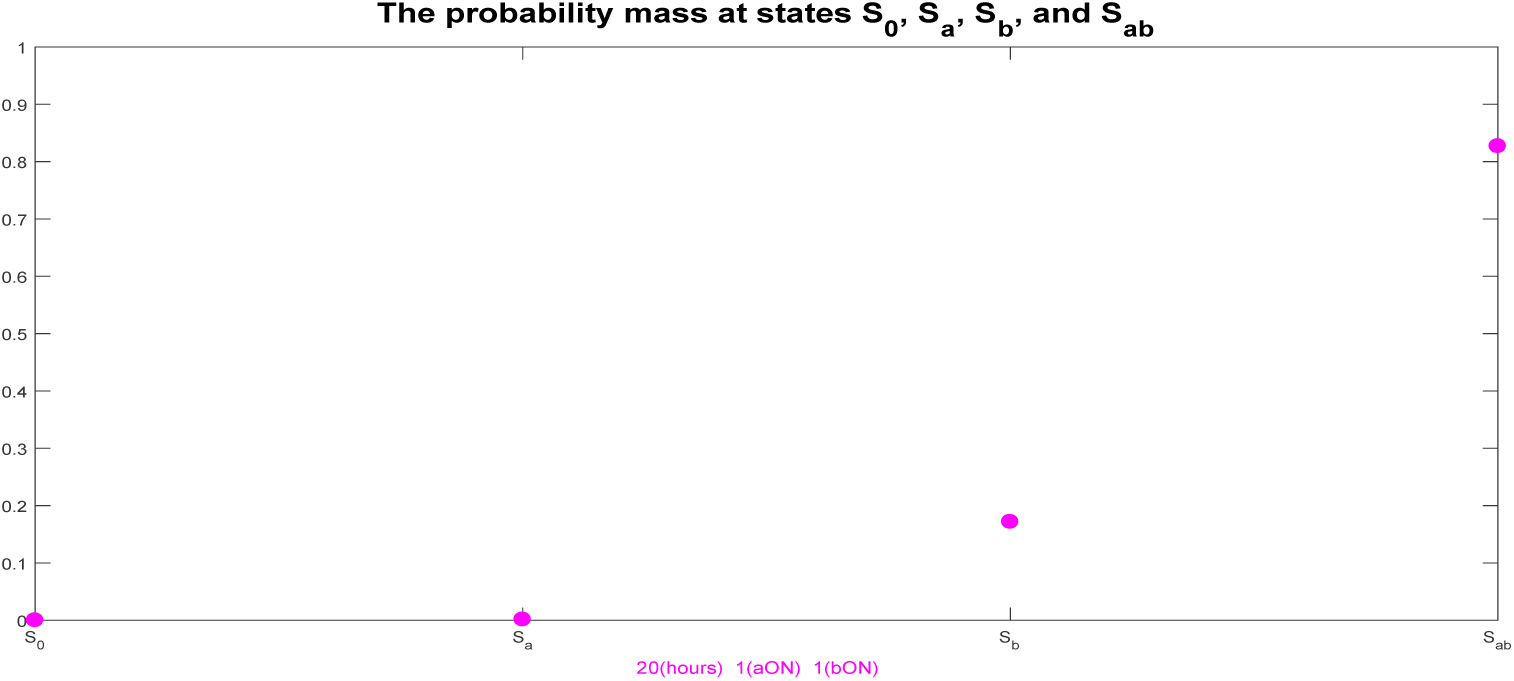
The temporal logic gate circuit has been running for 20 hours and it has reached stationary state. Input **a** has been turned on at 3 hours and input **b** has been turned on at 5 hours; subsequently, they were both on. The final population of cells is largely split between the cells that have recorded the “**a** then **b**” event and the cells that have only recorded the inducer **b** event.

## 3 Bayesian event detection and inference

### 3.1 The Bayesian framework

The Bayesian framework is a rigorous probabilistic method for representing uncertainty using probability distributions. This philosophy is rooted in probability as a logic [9, 18, 19, 20]. Within this framework, probability distributions are used to quantify uncertainty due to insufficient information, regardless of whether that information is believed to exist but is currently not available (epistemic uncertainty), or it is believed to not exist because of postulated inherent randomness (aleatory uncertainty). This makes the Bayesian framework appropriate for posing system identification problems, where postulated system models have parameters whose values are uncertain rather than random. Therefore, we view system identification as updating a probability distribution that represents our beliefs about models of a system based on new information from system response data.

One formulation of Bayesian system identification is given in [9], where *p* () describes a probably density function: Given observation data 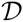 and a system model class 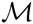 consisting of (a) a likelihood function 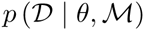, describing the plausibility of the data given a set of parameters, *θ*, and (b) a prior distribution, 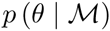, representing our initial beliefs about the relative plausibility of the possible values of the model parameter vector *θ*, find the posterior distribution 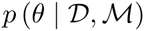 that represents our updated beliefs. For this, we employ Bayes’ Theorem:

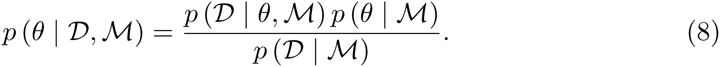

The likelihood function, 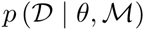, is the likelihood of observing the data *D* given that our forward model of the dynamical system has parameter values *θ*. The forward model in the Bayesian framework maps *θ* to a probability distribution on the outputs *y* (*t*). The normalizing factor in equation (8), 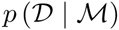, is the evidence for the model class 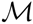. The evidence can be computed as

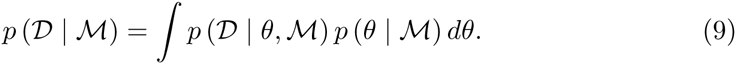

### 3.2 Constructing model classes from the forward models

In order to formulate the Bayesian inference problem for detecting events and determining the properties of these events, we create the following model classes. The first model class 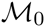 describes the dynamics of the cell population when there is no event, while 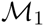 describes the dynamics of the population when there is an event described as the addition of chemical inducers **a** and **b**. The events in 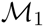 are parameterized by a vector *θ* defined in Table 4.

**Table 4:**
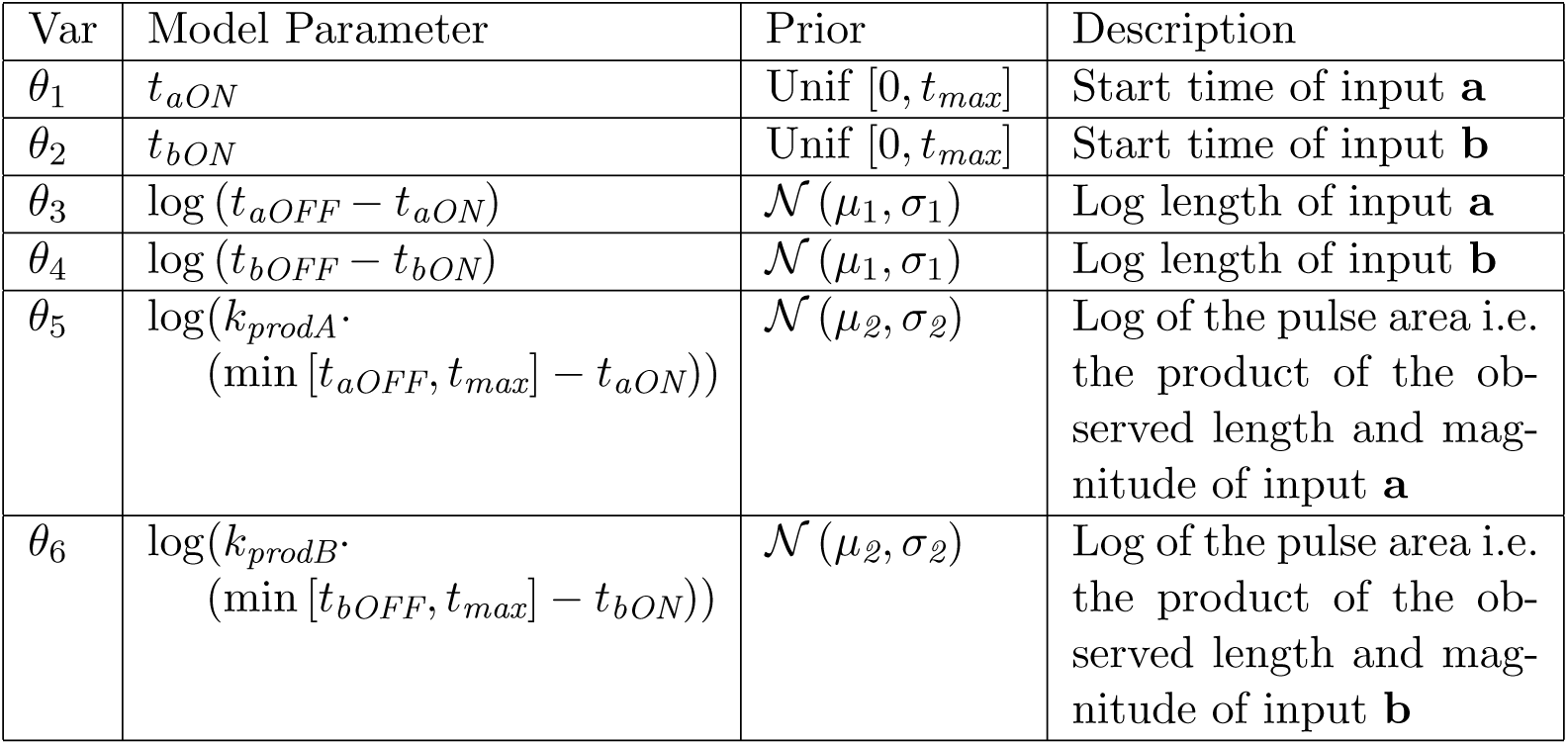
The variables, *θ*, parameterize the event that chemical inducers **a** and **b** are added based upon the start time, end time, and magnitude. We choose this parametrization to reduce the correlation between states and to avoid having prior distributions with bounded support, which can accelerate our sampling methods.

The forward models for 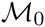 and 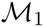 have the same structure, therefore we can construct the likelihood function for the model class in the same way. The forward model describes the evolution of a probability vector over DNA states, according to the CME. The data we are considering is the observed fraction of cells in a given DNA state at an instance in time. In order to find the likelihood of the data given our model, we consider two sources of uncertainty. First, in order to account for predictive modeling errors, we assume that the probability vector *p* (*t*), which we model as generating our observation *x* (*t*) at time *t* is not the same as the one generated by the forward model 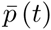, but comes from a distribution centered around the vector predicted by the forward model. Thus, in our model class, the forward model describes not the evolution of a probability vector, but the evolution of a distribution of probability vectors.

This distribution of probability vectors is modeled by a Dirichlet distribution with parametrization *α* (*t*), which we take to be 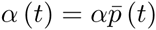. Here *α* is a constant that controls the variance of the “distribution of distributions”, i.e. the believed accuracy of our prediction. Secondly, we model the sampled number of cells in a given state, *x* (*t*) at time *t* using the multinomial distribution defined by the probability vector *p* (*t*) ∼ *Dir* (*α* (*t*)). The Dirichlet distribution is a common choice for quantifying uncertainty about a multinomial distribution since it is the conjugate prior of this distribution. By simultaneously considering these two sources of uncertainty, we are best able to replicate the uncertainty found in experimental data. When the number of measured cells is small, the sampling uncertainty will dominate, while when the number of measured cells is large, the prediction uncertainty will dominate. This formulation of the predictive model is summarized in Figure 8.

**Figure 8:**
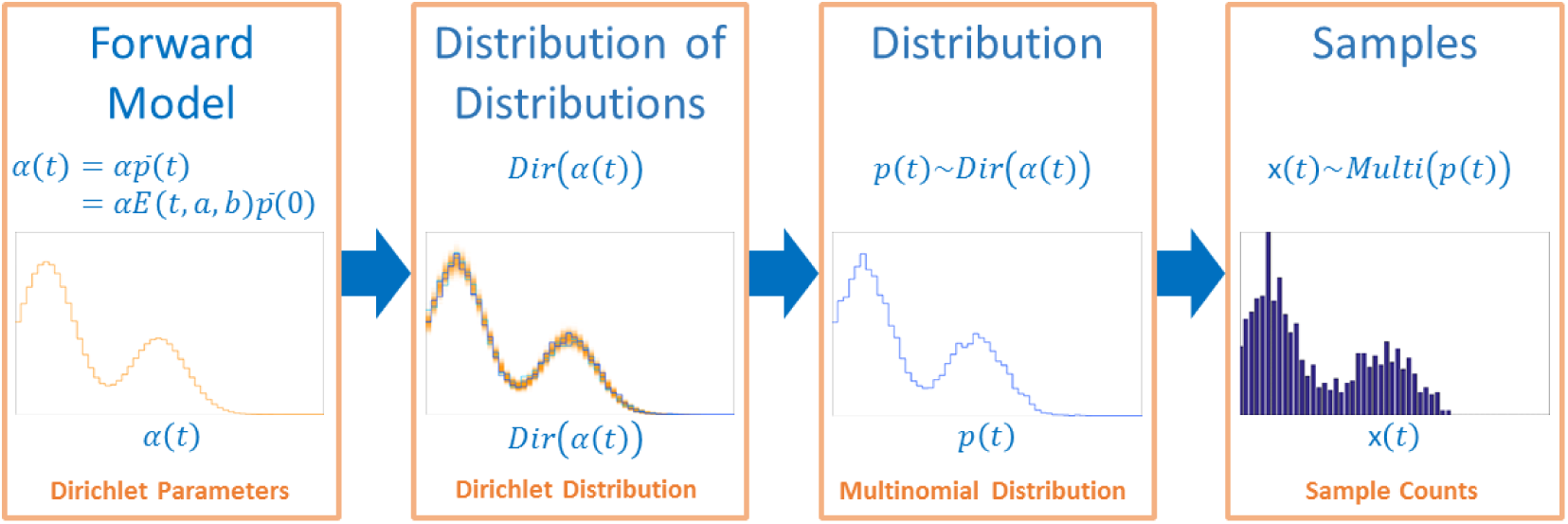
The probabilistic model used to construct the likelihood of the observed data is based on two sources of uncertainty: model prediction uncertainty and random sampling errors. We model the prediction uncertainty by having the CME evolve a Dirichlet “distribution of distributions” in time. Its mean is the probability vector for the standard CME evolution. Secondly, we model the random sampling error as drawing cells from a multinomial distribution chosen from our Dirichlet distribution.

Using our formulation of the model class, we can now define a likelihood function based on the forward model, Dirichlet distribution, and multinomial distribution as follows:

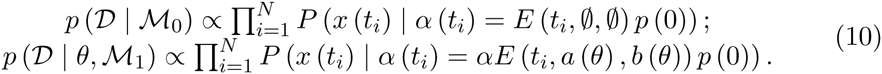

Here *x* (*t_i_*) are the cell counts at time *t_i_*, *α* (*t_i_*) is the parametrization of the Dirichlet distribution, and *N* the number of observations in time. This also models our prediction of cell population measurements as independent in time. We are only interested in likelihood functions up to a constant of proportionality for the computational methods we consider. Hence, equation (10) reduces to the following log likelihood function:

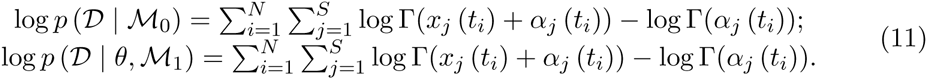

Here, *x_j_* (*t_i_*) is the number of cells in state *j* at time *t_i_*, *α_j_* (*t_i_*) is the parametrization of state *j* of the Dirichlet distribution at time *t_i_*, Γ is the Gamma function, and *S* is the total number of states in the state vector of the CME.

### 3.3 Detection and inference

Assuming that an event occurs, we can use Bayesian inference to infer the posterior distribution of the event parameters conditional on the measured data. We use the priors and likelihood functions in Section 3.2 to define the posterior distribution:

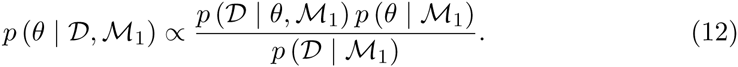

Using this function and Markov Chain Monte Carlo, we can then sample the posterior distribution as discussed in Section 3.4. We can also consider the probability of any event from 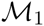 occuring given the cell population measurements, 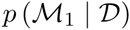. We assume that the prior probability of any event happening is known and defined as 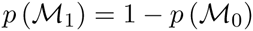. Therefore, we can perform event detection by computing

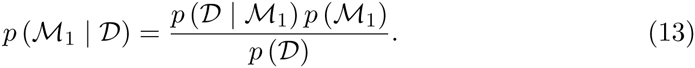

Using the law of total probability, 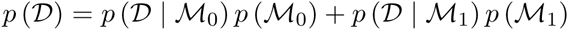, we find that

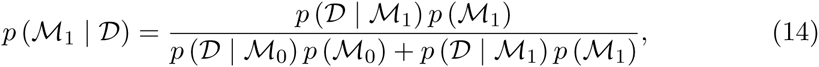

where 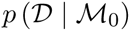 is defined in equation (10). Hence, it remains to estimate the evidence, 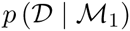, as:

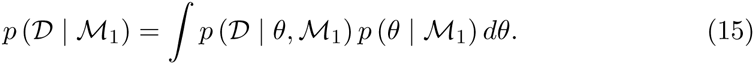

Estimating this quantity is difficult, but several computational methods have been introduced to provide good estimates for model class selection, as we discuss in Section 3.4.2.

### 3.4 Computational methods: MCMC

Sampling methods are typically used to solve Bayesian inference problems. We can estimate quantities with respect to the posterior distribution using a population of samples as follows:

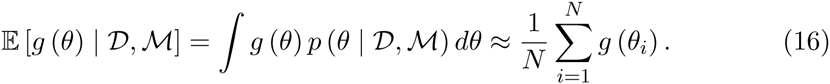

The most common family of sampling methods for Bayesian inference is Markov Chain Monte Carlo (MCMC) [6]. In MCMC, we create a Markov chain defined by a transition rule or kernel and whose stationary distribution is the desired posterior. In order to have accurate estimates, the samples must discretely capture the posterior distribution in a probabilistically appropriate way. By the Markov chain central limit theorem, we can estimate the quality for the mean estimate of a finite-variance stochastic variable based on the number of samples and the correlation function of the Markov chain. When selecting and implementing a MCMC method, we seek to minimize the correlation between the states of the chain. In this way, we decrease our estimate variance and we also minimize the time it takes for chain to reach its stationary distribution.

#### 3.4.1 MCMC implementations

We consider two basic MCMC implementations for sampling the posterior of *θ* for 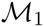: Metropolis-Hastings and Adaptive Metropolis-Hastings. Metropolis-Hastings produces a Markov chain with a desired stationary distribution *π* (*θ*) by designing a transition kernel *K* (*θ^′^* | *θ*) such that the Markov chain is ergodic and reversible [21, 22]. Reversibility is a sufficient condition for the existence of a stationary distribution. Reversibility holds under the detailed-balance condition

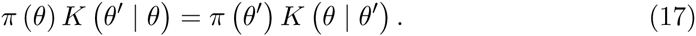

This means that we can choose any transition kernel *K* (*θ^′^* |*θ*) and maintain the stationary distribution *π* (*θ*), as long as the condition in equation (17) holds. For any proposal distribution *Q* (*θ^′^* |*θ*) such that *Q* (*θ^′^* |*θ*) ≠ 0 if and only if *Q* (*θ | θ^′^*) ≠ 0, we can construct such a *K* (*θ^′^* |*θ*) by proposing a candidate sample *θ^′^* according to *Q* (*θ^′^* |*θ*). Then we accept the candidate *θ^′^* with probability *α* from equation (18). If the candidate is rejected, the current sample *θ* is repeated:

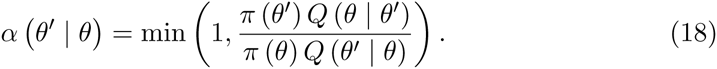

This leads to the Metropolis-Hastings algorithm:

1. Initialize the state *θ* _1_ randomly, usually according to the prior; set *n* = 1.
2. Pick a candidate state 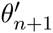 according to the proposal 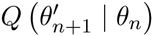.
3. Accept or reject the candidate according to a sampled uniform variable *ζ* on [0, 1]:

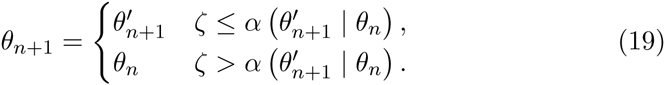
4. Increment *n* and go to step 2.

We choose the proposal distribution to be a Gaussian distribution. Typically, several runs are used to tune this distribution such that good performance is achieved. When the posterior distribution is a Gaussian, the optimal proposal distribution is a Gaussian with covariance 2.38^2^Σ*=d* where *d* is the dimension and Σ is the covariance of the posterior [6].

Adaptive Metropolis-Hastings has the same structure as the Metropolis-Hastings; however, the proposal distribution is adapted over time in a way that still maintains the stationary distribution of the Markov Chain.

#### 3.4.2 Estimating the evidence

Estimating the evidence for the model class defined in equation (15) can be quite challenging since it requires solving a difficult high-dimensional integral. Many approximation methods have been developed to estimate this high-dimensional integral such as the Laplace Approximation, importance sampling, and multilevel methods like TMCMC [9, 23]. When the prior and posterior distributions can be well approximated using a Gaussian, we can use the following approximation to estimate the evidence:

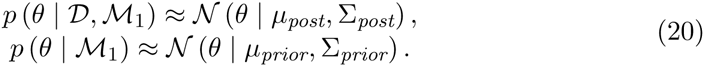

By replacing the posterior and prior in Bayes’ Theorem by these approximations and rearranging terms, we find that

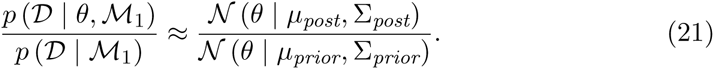

Thus, by taking the log and then taking the mean over our samples, we can approximate the log evidence as

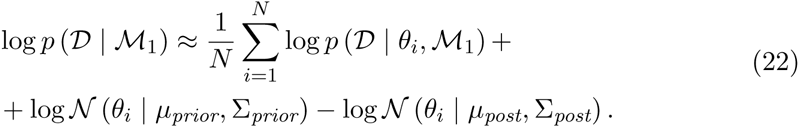

## 4 Applying the Bayesian approach to temporal logic gate data

In order to judge the efficacy of our Bayesian framework, we test it over a set of chemical inducer pulse properties. For these tests, we use the same setup, but vary the length of the inducer pulses. In all cases, inducer **a** is added at 3.0 hours and inducer **b** is added at 5.0 hours. Each inducer has a production rate of 0.5 (*μ*^3^hr)^−1^. For the Bayesian inference problem, the log pulse duration prior is 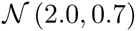 (2.0; 0.7) and the log pulse area prior is 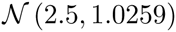 (2.5; 1.0259). The forward computations of the matrix exponential vector products uses Expokit [24]. In both cases, the estimated posterior probability of any event using the Gaussian approximation was 1.0.

### 4.1 Case 1: Nominal pulse duration

First, we consider the parameters for a typical event that has inducer **a** on for 7.0 hours and inducer **b** on for 5.0 hours. We use an adaptive MCMC method to generate posterior samples. The evolution of the Markov chain is illustrated in Figure 9. The adaptation is separated into several periods to effectively trade off between parameter values that minimize both the burn-in period and the correlation. The chain starts by using a wide fixed proposal distribution for the first 400 iterations. Then it switches to a fixed narrower proposal for the next 600 iterations. It then uses an adaptive method for the next 500 iterations and then restarts the adaptive method for the remainder of the 5000 iterations. By breaking up the iterations, we can avoid having too much memory in the process and ensure that the adaptive MCMC is just learning from samples after burn-in. We consider samples after the 2000^*th*^ iteration to belong to the posterior. Since MCMC produces correlated samples and our acceptance rate is near the optimal of 0.25, our efficiency is about 0.025 independent samples per iteration after burn-in.

**Figure 9:**
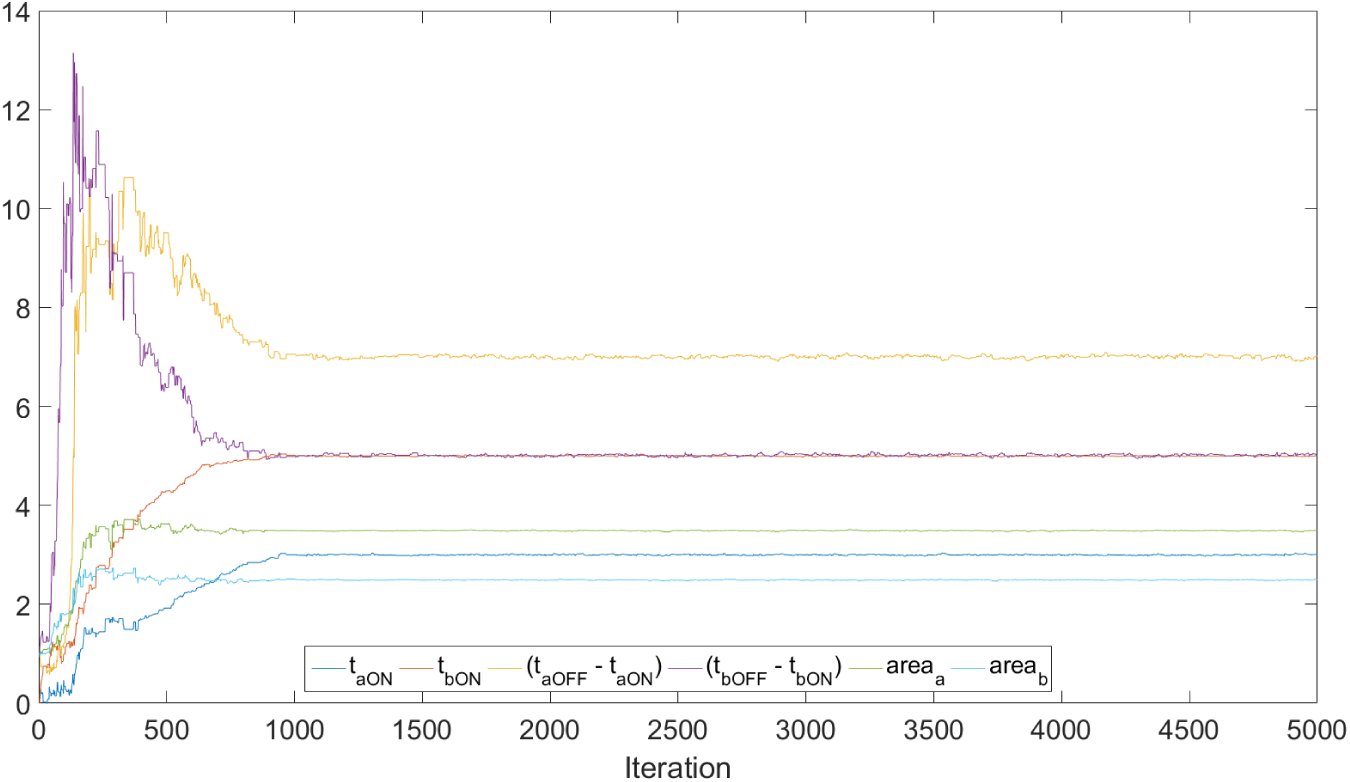
Evolution of the Markov Chain using Adaptive MCMC for Case 1. We can see that the Markov Chain successfully finds and samples the posterior.

In Figure 10, we plot the histogram for the posterior samples and the correlation diagrams for each of the parameters. While there is still some correlation between the pulse length and area, our choice of using pulse area instead of pulse amplitude does eliminate much of the dependence. However, there is significant correlation between the start time of the pulse and the length of the pulse. From these plots, we find that all the posterior samples histograms are globally identifiable since they are unimodal. The mean and standard deviation estimates of the posterior distribution are found in Table 5, where there is very good agreement between the mean estimates and true values.

**Figure 10:**
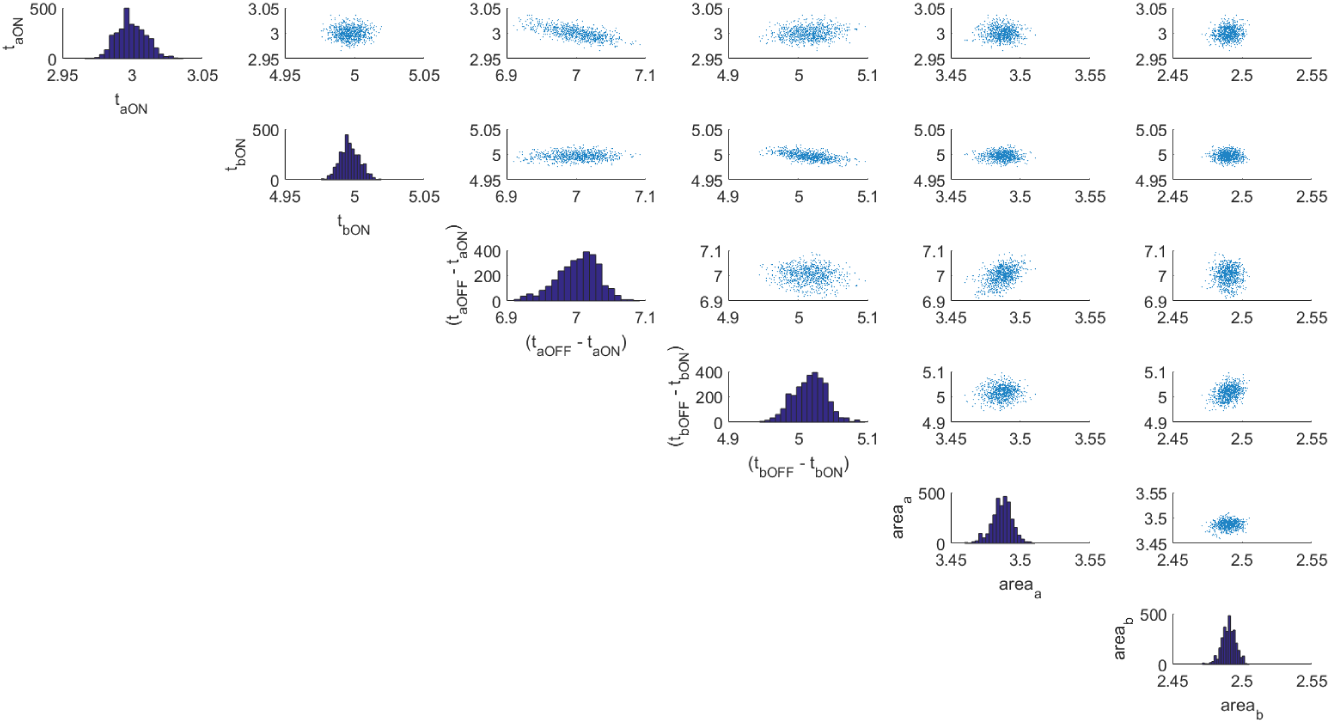
Histograms of the posterior sample and scatter plot showing the correlation in the posterior for Case 1.

**Table 5:**
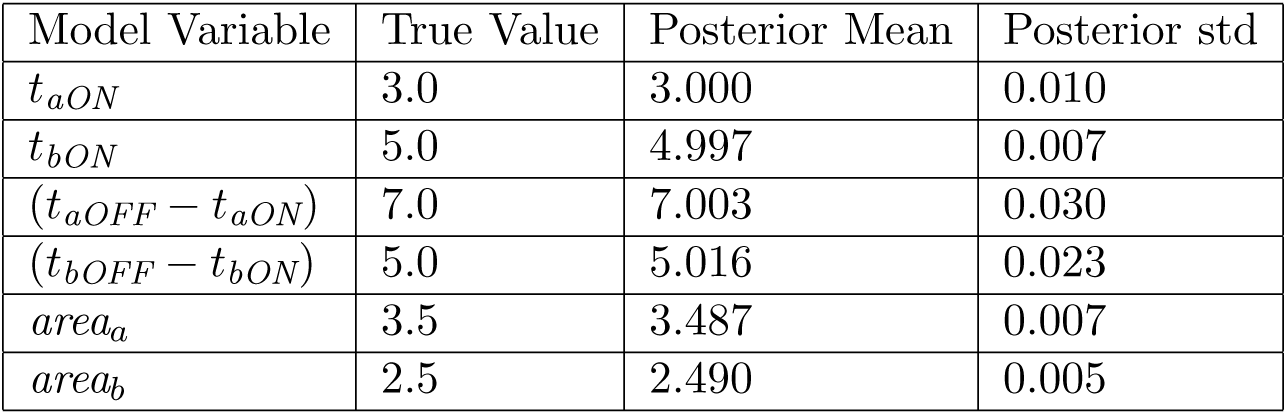
Posterior estimates for Case 1

### 4.2 Case 2: Short pulse of inducer b

Similarly, we can consider a more challenging case when the pulse of inducer **b** is much shorter, only 0.5 hours long, but everything else is the same. We can see the posterior histograms and correlation diagrams in Figure 11 for 2000 posterior samples. We see a similar correlation structure as that observed in Case 1. The posterior is still globally identifiable, making it a good candidate for the approximate solution to the event detection problem. The posterior values in Table 6 are also in good agreement with the true values. We can see that compared to Case 1, the posterior for this example has a larger coefficient of variation for the inducer **b** parameters, indicating that it is less identifiable.

**Figure 11:**
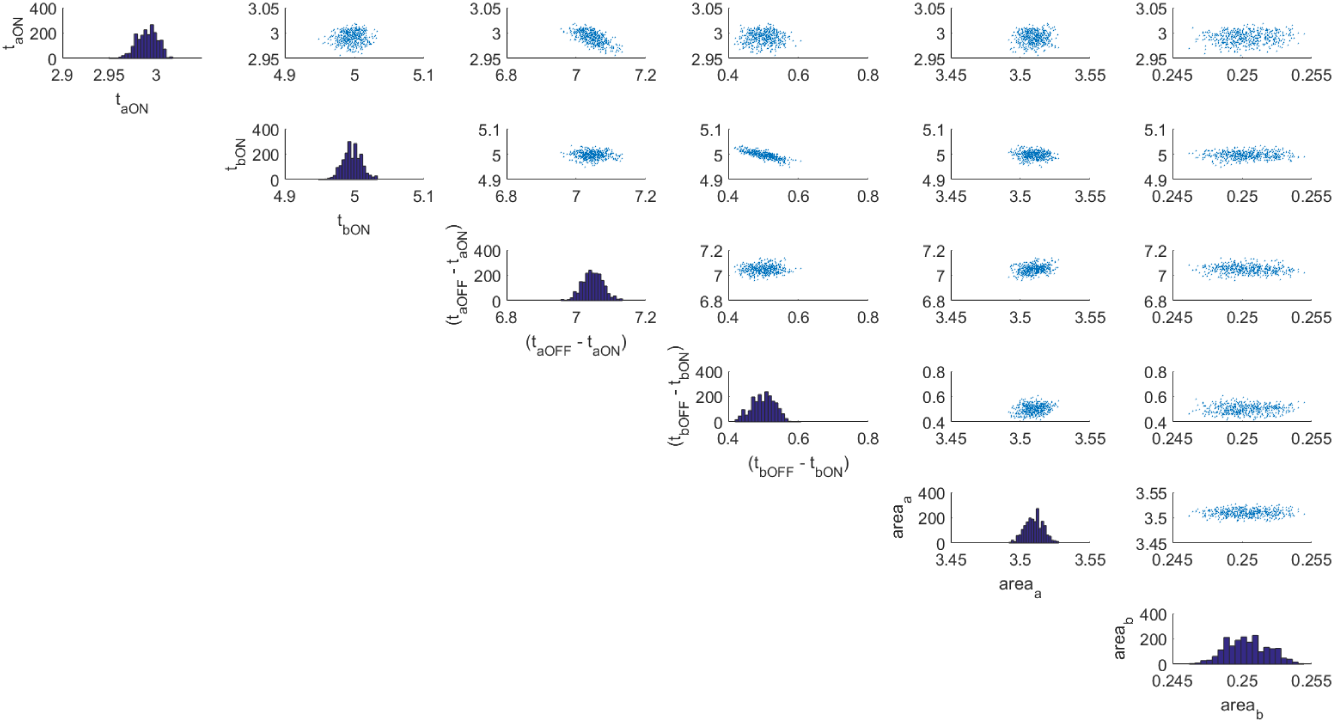
Histograms of the posterior sample and scatter plot showing the correlation in the posterior for Case 2.

**Table 6:**
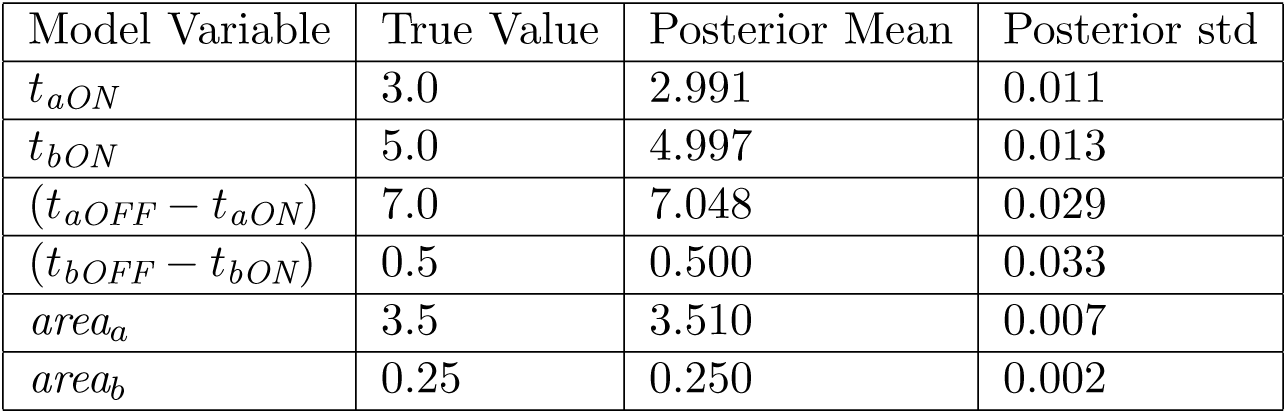
Posterior estimates for Case 2

### 4.3 Case 3: Undetectable inducer pulses

Undetectable events occur when the length of the chemical inducer pulse is smaller than 5 minutes, the pulse amplitude is very low, or the modeling error is very high. Clearly, a pulse shorter than the time interval at which we take measurements will not be detected. A low amplitude pulse is also undetected when *k_leakA_ ≈ k_prodA_* or *k_leakB_ ≈ k_prodB_*, which are not biologically relevant scenarios. The last case of undetectable events is the most realistic for biological systems. It could be the case that our model was a very poor representation of the true biological system that produced the pulses. Then the variance of the noise in our “distribution of distributions” could be 100 times larger than the current acceptable noise level and then the pulses would be undetectable. However, we know from [3] that the model that has been validated against experimental results to prevent this scenario.

## 5 Conclusion

Using the framework of Bayesian inference, we have answered questions about events recorded in the heterogeneous distributional response of a cell population. We were able to identify the occurrence, timing, and amplitude of chemical inducer pulses in a temporal logic gate circuit. We used the cell population response to determine whether an event of chemical induction of integrase expression had occurred. We also obtained accurate results for chemical inducer pulse timing, length, and amplitude using Markov Chain Monte Carlo methods. We detected and identified chemical inducer pulses as short as half an hour, as well as all pulse amplitudes that fell under biologically relevant conditions.

In future work, we plan to implement the Bayesian inference of chemical inducer properties using Transitional Markov chain Monte Carlo (TMCMC) [23]. TMCMC will more accurately find and sample complicated distributions that arise when the model is close to unidentifiable. This can occur when there are short chemical inducer pulses and when the model poorly captures the experimental cell population behavior. TMCMC will enable us to not rely on Gaussian assumptions for evaluating the evidence in equation (9) to distinguish between chemical inducer pulses being on or off. Due to better computational performance than MCMC or Adaptive MCMC, we aim to increase the complexity and the number of parameters that describe our chemical inputs. In the future, we aim to solve the inference problem for a chemical inducer class of functions that correspond to square wave trains, which represent inducers repeatedly turning on and off.

